# Sampling for disease absence—deriving informed monitoring from epidemic traits

**DOI:** 10.1101/443895

**Authors:** Yoann Bourhis, Timothy R. Gottwald, Francisco J. Lopez-Ruiz, Sujin Patarapuwadol, Frank van den Bosch

## Abstract

Monitoring for disease requires subsets of the host population to be sampled and tested for the pathogen. If all the samples return healthy, what are the chances the disease was present but missed? In this paper, we developed a statistical approach to solve this problem considering the fundamental property of infectious diseases: their growing incidence in the host population. The model gives an estimate of the incidence probability density as a function of the sampling effort, and can be reversed to derive adequate monitoring patterns ensuring a given maximum incidence in the population. We then present an approximation of this model, providing a simple rule of thumb for practitioners. The approximation is shown to be accurate for a sample size larger than 20, and we demonstrate its use by applying it to three plant pathogens: citrus canker, bacterial blight and grey mould.

## Introduction

When it comes to disease management, surveillance programs have two different objectives: establishing disease absence in host populations, or ensuring an early detection of any disease outbreak (Parnell et al., 2017). Early detection is essential to disease control mitigation, timely reactions generally being more successful and less detrimental for the host population (Cunniffe et al., 2016). For example, Carpenter et al. (2011) showed for foot-and-mouth disease, that when delaying the detection from 7 to 22 days after the initial infection, the containment measures required the culling 30 times more host animals. Likewise, surveillance programs are operated to establish the absence or presence of emerging strains of endemic pathogens, hence enabling trade certifications for instance. Examples of these are emerging strains of plant pathogens that are insensitive to the fungicides applied to control them, or strains that are virulent^1^ in a crop cultivar by having resistance breaking genes.

Monitoring a disease requires the assessment of the pathological status of sampling units. Assessments generally occur at the level of the host individual (*e.g.* for ash dieback, Woodward & Boa, 2013), but sometimes for convenience the sampling unit is a subpopulation like a farm (*e.g.* for foot-and-mouth disease, Keeling et al., 2001), or a field (*e.g.* for bacterial blight in rice, Koyshibayev & Muminjanov, 2016). In any case, when the samples all return negative, it is still very important to account for the chance that the pathogen was present but undetected (Cannon, 2002). Therefore, declaring disease absence is then a probabilistic evaluation, more samples making it less likely the pathogen was missed.

The incidence^2^ of a pathogen, noted *q* hereafter, is the proportion of the host population infected. Estimating the incidence from a sample where all assessments return negative for the pathogen can be defined as a zero-numerator problem, *i.e.* estimating the probability of an event from data in which it has not occurred yet (Hanley & Lippman-Hand, 1983; Winkler et al., 2002). We thus want to calculate the probability density, *p*(*q| notfound*), of the incidence *q* given that none of the sampling units is assessed as infected. This can be done by deriving *p*(*notfound |q*) from the exponential distribution, and then reversing it according to Bayes’ rule, assuming a uniform prior *p*(*q*). A practically useful quantity is the incidence *q_X_* for which

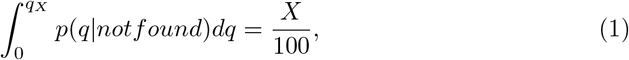

thus giving the upper bound of the *X*% confidence interval of *q*. This upper limit gives the highest, still likely, incidence given a sampling effort. The rule of three is a very common rule of thumb to estimate the upper limit of the 95% confidence interval (Louis, 1981). For example, considering we have a random sample of size *N*, all returning negative, the upper limit can be approximated by 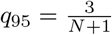 (Hanley & Lippman-Hand, 1983). This very practical rule of thumb can be used to identify a sampling effort *N* that can ensure that infection is below a threshold value.

To ensure pathogen absence from an area over an extended time interval, the host population have to be sampled repeatedly. Incidence estimation should then account for the change of incidence between the rounds caused by the epidemic dynamics. In this regard, Metz et al. (1983) accounted for the time dependence of samples due to the epidemic dynamics when they assessed the level of epidemic risk associated with a given sampling effort. However, when it comes to the incidence estimation problem, the epidemic temporal dynamics is neglected while the focus is more likely set on the spatial dependences of the samples due to the epidemic spread (Cameron & Baldock, 1998; Cannon, 2002; Coulston et al., 2008).

Accounting for the epidemic dynamics, we address the incidence estimation problem in the case of *disease absence* sampling (as Parnell et al., 2012, did with *first discovery*). We present a model estimating the pathogen incidence in a population, being given a sampling effort and an epidemic growth rate. We then derive an approximation of this model (in the way of Alonso Chavez et al., 2016) providing a practical and simple way to derive information from a negative sampling. This epidemically informed approximation proves itself accurate and flexible enough to account for the asymptomatic period of the disease. Finally, we apply this model to three case examples: citrus canker in an orange orchard, the invasion of virulent pathogen strains of bacterial blight of rice and the invasion of fungicide resistant pathogen strains in grey mould of grape.

## Materials and methods

A monitoring program typically consists of batches of *N* samples randomly or regularly distributed in space, and regularly iterated over *K* monitoring rounds with time intervals of ∆ time units. Parnell et al. (2012) has shown how to use this particular structure to derive the pathogen incidence when first detection occurs. In this section we use a similar method to estimate the incidence when no infected sample has returned.

### One monitoring round

Considering an incidence *q* in a population, the probability for a sample to return negative is given by *p*(*notf ound |q*) = 1 – *q*. A sample of size *N* will therefore return entirely negative with probability:

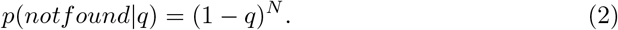

Now, given a monitoring round returned negative, what does it tell us about *q* ? We can derive *p*(*q| notf ound*), the incidence given no detection, from *p*(*notf ound| q*) using the Bayes’ theorem. Bayes’ theorem relates those probabilities by:

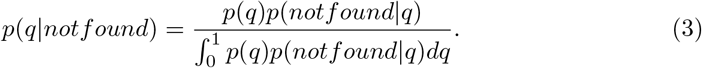

The value *p*(*q*) is the prior probability density of *q*. Assuming no information on *q*, we set *p*(*q*) to be a uniform and uninformative prior. Substituting Eq. 2 into Eq. 3 then gives

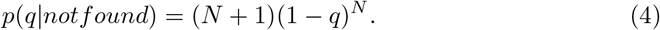

As mentioned in the introduction, of particular interest is the upper limit of the *X*% confidence interval given by Eq. 1 which, for the exponential distribution given by Eq. 4, gives

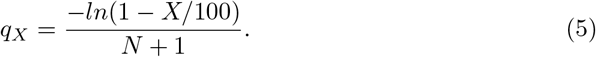

Put in words, if we set a maximum incidence *q_X_* below which we are satisfied to consider the host population to be free of disease, we can derive a sampling effort *N*, that will ensure *X*% of the undetected diseases to have incidences smaller than *q_X_*.

### Two monitoring rounds

Having two monitoring rounds can be seen as increasing the size of the sample:

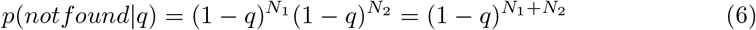

where *N*_2_ is the size of the current monitoring round and *N*_1_ is the size of the previous one. In this equation, the sizes of the historic and recent monitoring rounds are powers of the same probability of negative sampling: 1 − *q*. However, as mentioned in the introduction, the incidence of an infectious pathogen increases through time. Therefore, non-detection in the last monitoring round occurred over a larger incidence than the previous one.

As we are focused on absence sampling, we are interested in epidemics with low incidences, so *q*≪ 1. It is well established that at low incidence epidemics grow exponentially (van der Plank, 1963; Faria et al., 2014; Bartlett et al., 2016). We thus assume *q*(*t*) = *q*(0)*e^rt^* where *t* is the time and *r* is the epidemic growth rate. In the time interval ∆ between two monitoring rounds the pathogen incidence has grown by a factor *λ* = *e^r^*^∆^, or another words, the incidence in monitoring round *i* was a factor λ^*−*1^ smaller than the incidence in monitoring round *i* + 1.

For our two-round case we thus have

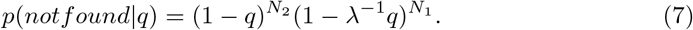

### K monitoring rounds

Building on this epidemic model, we can now generalise Eq. 7 to *K* monitoring rounds:

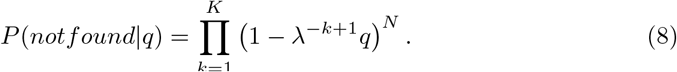

We can use the Bayes’ theorem to compute the probability density of the incidence given non-detection after *K* monitoring rounds as:

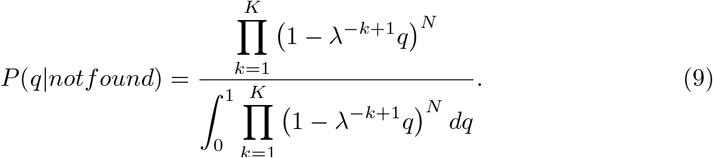

### An approximation

Computing the probability density given by Eq. 9, as well as the subsequent upper limit *q_X_*, requires integrations which need to be approximated numerically. It is computationally inexpensive but still requires a computer program to be used. Here we develop an approximation which makes the computation of *p*(*q| notf ound*) simple enough to be useful for practitioners. It gives a rule of thumb in planning a monitoring program for a given disease.

The approximation is built on the following two assumptions: (1) the sampling size *N* is large enough (*N >* 10), and (2) the incidence *q* is small. Both assumption are realistic as 10 is a relatively small sampling size, and as we are interested only in cases with very low incidence. Using our assumption that *q* ≪1, we can approximate (1 *q*)^*N*^by *e^−qN^*. Substituting this in Eq. (3) results in:

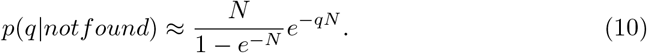

And following our assumption that *N >* 10, this equation can be approximated by

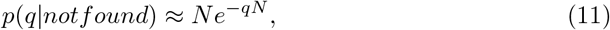

Plugging Eq. (11) into Eq. (1) results in

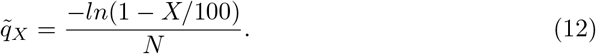

Similarly, for two monitoring rounds we find

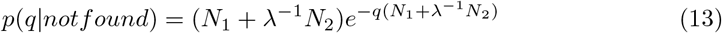

and

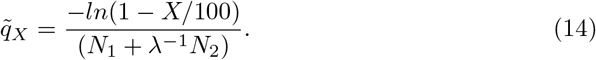

If now we generalise to *K* monitoring rounds, it results in

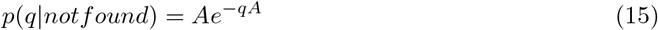

and

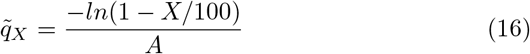

where A is given by:

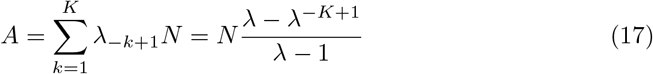

Before going further using the approximation, its accuracy needs to be evaluated. First, the approximated and the exact probability densities *p*(*q| notf ound*) are compared visually. And then, we investigate more carefully the difference between their respective confidence intervals *q_X_* and *q*_*X*_. The comparison is likely to depend on the sampling effort as well as on the epidemic growth rate, therefore the accuracy is evaluated for wide ranges of the relevant parameters.

## Results

### The exact model

Figures 1 clearly shows the effect of an epidemic increase (*i.e. λ >* 1) as compared to a situation, as previously published (see *e.g.* Cameron & Baldock, 1998; Cannon, 2002, for absence sampling), where incidence *q* is assumed constant over time (*i.e. λ* = 1). Figures 1 exposes that using the classical rule of 3 for a monitoring program extended in time would result in significant underestimations of *q*_95_, as it would for any confidence level. The severity of these underestimations increases with the epidemic growth rate and the time interval between rounds. As expected, the upper bound *q_X_* of the confidence interval for *q* decreases with increasing sample size *N* and increasing number of sampling rounds *K*. The faster the epidemic grows, the larger *λ*, and the larger *q_X_*, which is also to be expected. What is less obvious but interesting to note is that if we compare monitoring programs with the same sampling effort *N ⋅ K* (lower left versus upper right panels in Figure 1), we see that *q_X_* is lower for monitoring programs that are shorter in time (smaller *K*). This finding is consistent for other parameter values. However in reality we do require a monitoring program to extend over long period of time to ensure pathogen absence for the entire period.

**Figure 1.**
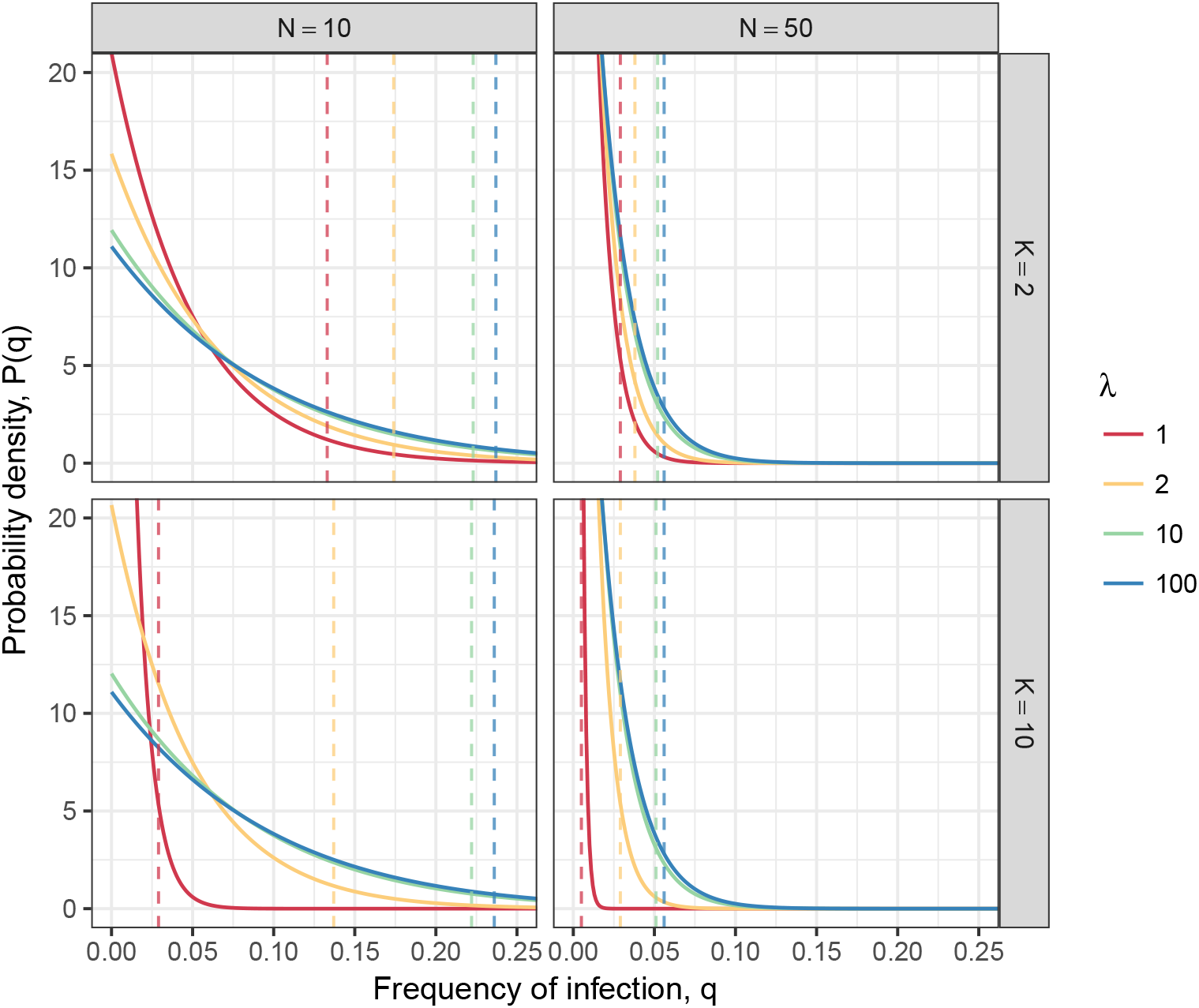
Probability density of the incidence given by Eq. 9 depending on the sampling effort (size *N* and repetition *K*) and on *λ* = *e^r^*^∆^, the factor by witch the epidemic grows between two sampling rounds. The dashed lines are the upper bounds *q_X_* the 95% confidence interval.

The impact of *λ* on the incidence can be decomposed to investigate the impact of the growth rate *r* and the time interval between rounds ∆. Since they are defined by *λ* = *e^r^*^∆^, they have the same impact on disease incidence, which is illustrated by the diagonal symmetry in Figure 2. This picture focus only on *q_X_* instead of the whole probability density. Figure 2 also delineates a plateau for large values of *λ*, above which a faster epidemic growth, or a larger monitoring time interval, does not significantly increase the incidence of the undetected pathogen. This is also visible in Figure 1 where the probability densities for *λ* = 10 and *λ* = 100 are very similar, despite the order of magnitude change in *λ*. Finally, Figure 2 illustrates the greater impact of the sample size *N* than the number of rounds *K* on the epidemic risk.

**Figure 2.**
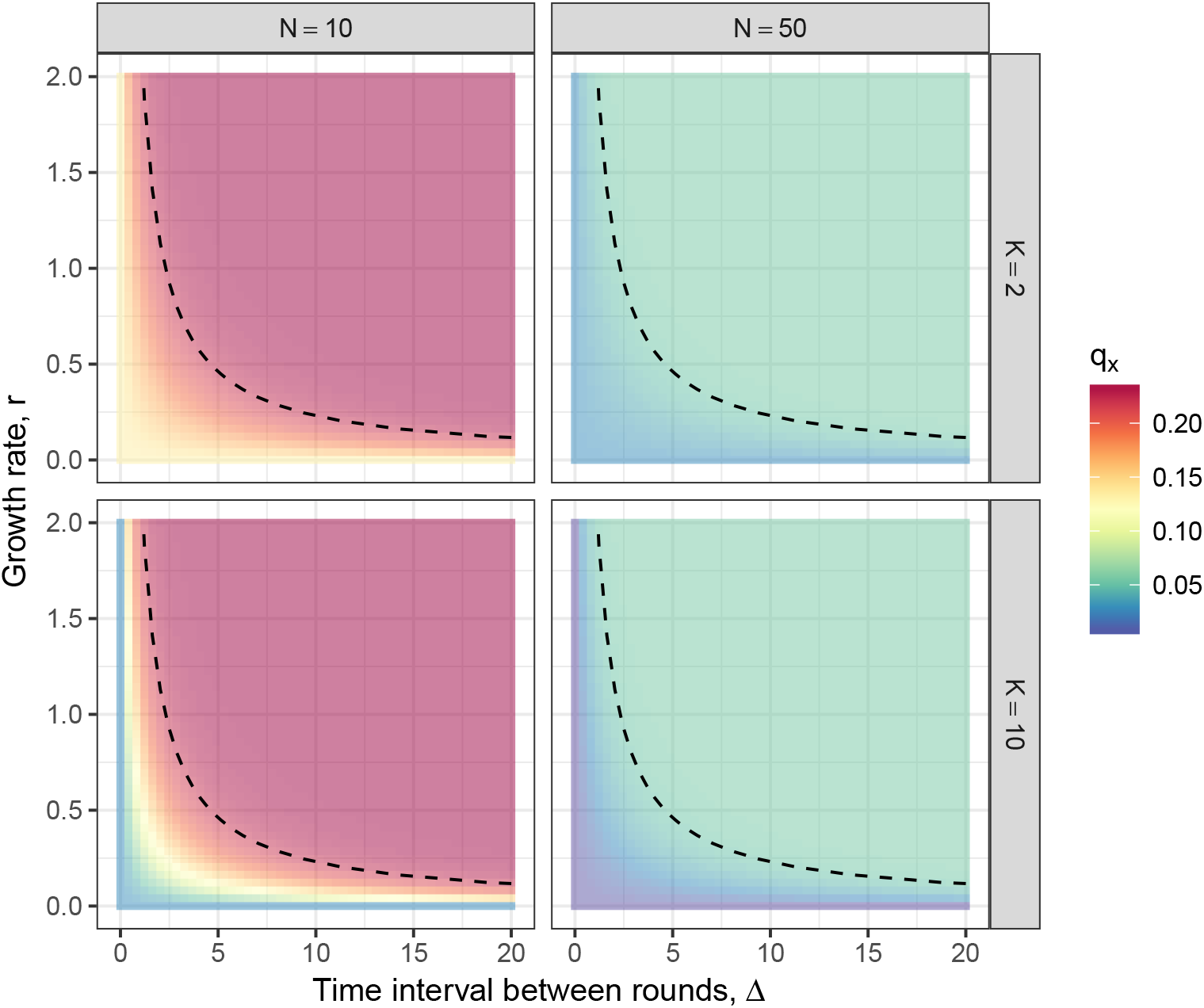
Upper limits of the 95% confidence intervals, *q*_95_, as a function of the two components of *λ*, *i.e.* the time interval between rounds ∆ and the epidemic growth rate *r*. The four panels show four sampling efforts defined by *K* and *N*. The dashed black lines figure the *λ* = 10.

### Accuracy of the approximation

In developing an approximation, our aims are twofold: (1) provide an equation featuring the model behaviours described in the previous subsection, and (2) provide an equation simple enough that it can be solved “on the back of an envelop” when designing a monitoring program. Figure 3 compares, for the case with two monitoring rounds (*K* = 2), the exact and approximated probability densities (respectively given by Eqs. 9 and 15). We see that the exact and approximated density curves are barely distinguishable whatever the sampling size and epidemic speed. However, if we take a closer look at our index of interest *q_X_*, we see that low values of *N* cause significant inaccuracy (figured by the shaded areas). This illustrates why the approximation does not hold for *N <* 10. Figure 3 also shows a tendency towards inaccuracy when the epidemic growth or the monitoring intervals increase.

**Figure 3.**
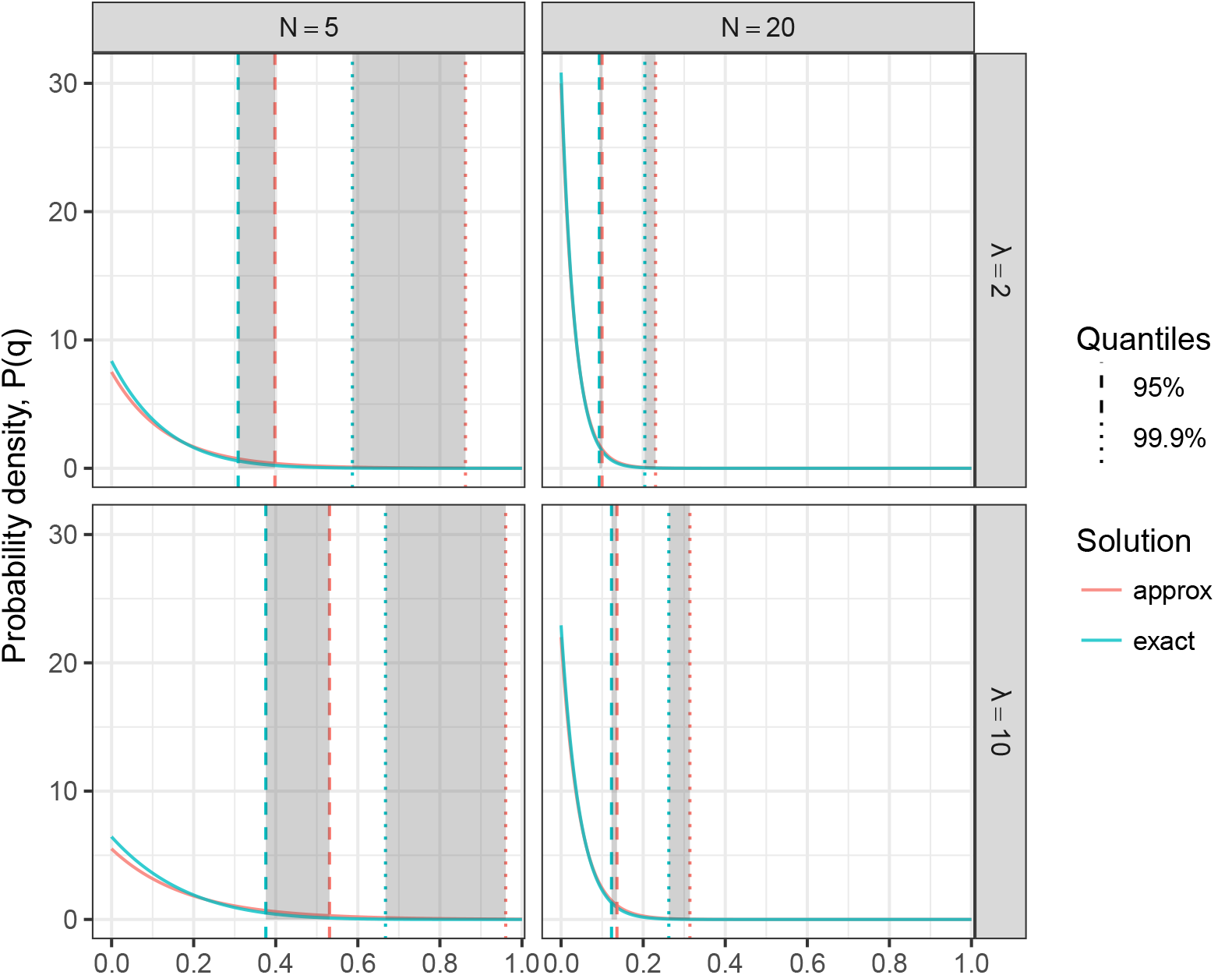
Exact and approximated probability densities of the incidence *p*(*q| notf ound*) for two monitoring rounds. The vertical lines show the 0.95 and 0.999 quantiles (indicating the upper bounds of the respective confidence intervals). The exact *q_X_* is derived from the numerical integration of Eq. 9, while the approximated *q*_*X*_is given by Eq. 15.

The effect of *K* (the number of sampling rounds) is better visualised if we focus on the relative error between the approximated and exact upper bounds 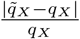. Figure 4 confirms the trends previously observed: the accuracy increases with the sampling size and decreases with the epidemic growth rate and time interval between samples. We see that past *N* ≈ 20, good levels of accuracy of the approximation is achieved, even for large epidemic growth rates. It also seems that going from *K* = 20 to *K* = 100 does not improve the accuracy and we reach a plateau. Finally, the accuracy increases with the likeliness of the event under study (here from 0.1% in bottom row to 5% for top row), which is also illustrated by the shaded areas in Figure 3.

**Figure 4.**
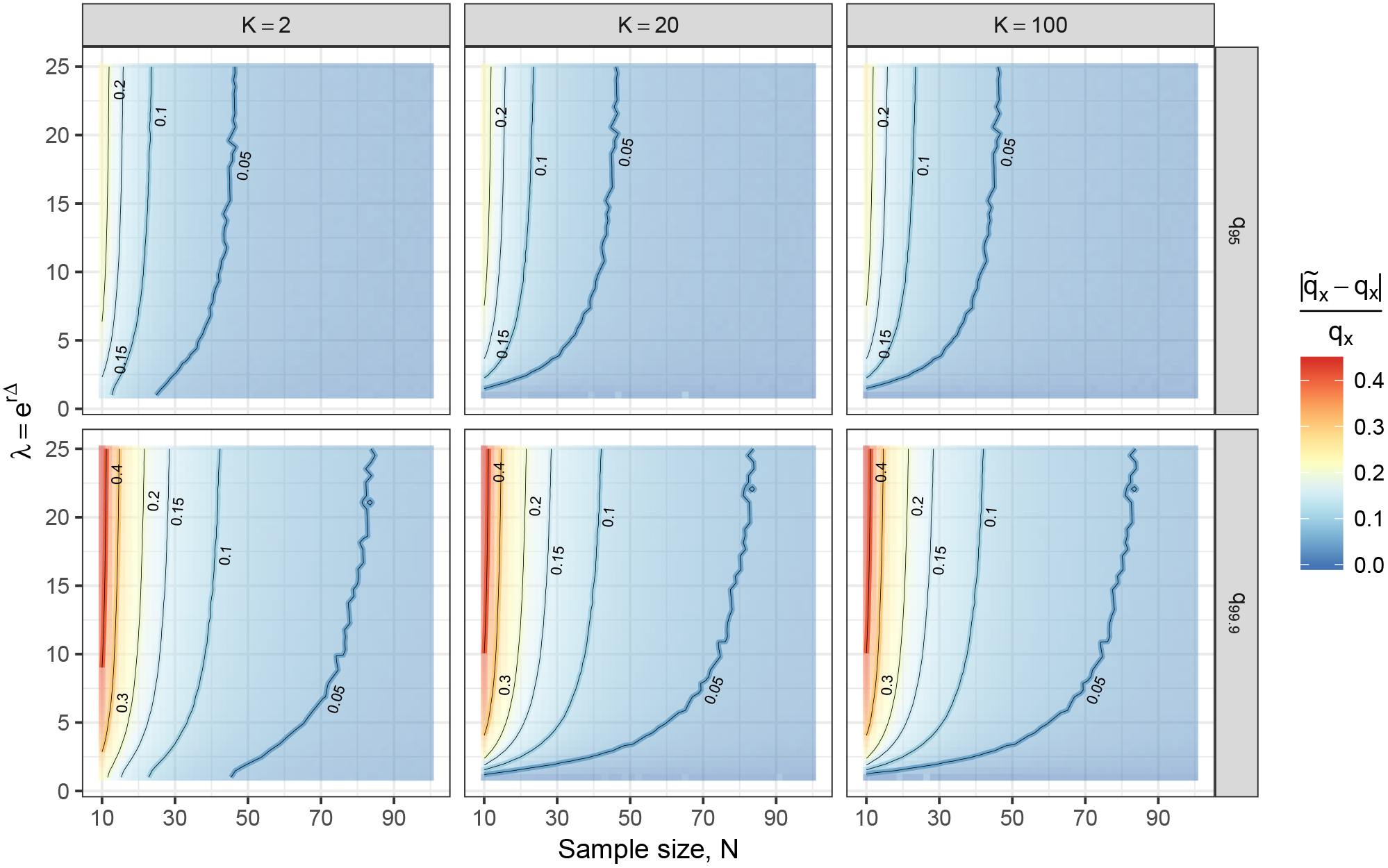
Relative difference between quantiles (for 2 levels of confidence, *i.e. q*_95_ and *q*_99.9_) of the exact and the approximated incidence (*q* and *q*) depending on *N*, *K* and *λ*. The contour lines and the background colours quantify the error.

## Applications

Having established that our approximation is accurate for sampling sizes *N >* 20, we turn towards three applications of the model. However before this is possible we need to discuss the asymptomatic period characteristic for most pathogens.

### Accounting for an asymptomatic period

After infection, the host is not detectable for a duration of time that depends on the pathogen species. This asymptomatic period is longer for visual assessment than for molecular diagnostics, but exists for each assessment method. It corresponds to the time needed by the host to develop detectable symptoms, *i.e.* outreaching a detection threshold. Since we need to estimate the possible incidence of all infected hosts, and not only of hosts with detectable infection, we need to take this asymptomatic period into account.

During the asymptomatic period, the newly infected hosts, that are not yet detectable as such, can still spread the pathogen. Therefore their impact on epidemics can be considerable, especially in the early stage of the disease as illustrated by Figure 5. Because of the exponential dynamics of the early epidemics, the difference between what we can observe, *i.e.* the detectable incidence *q*, and what is actually spreading the pathogen, *i.e.* the total incidence *q_T_*, promptly becomes significant even for fairly short asymptomatic periods.

**Figure 5.**
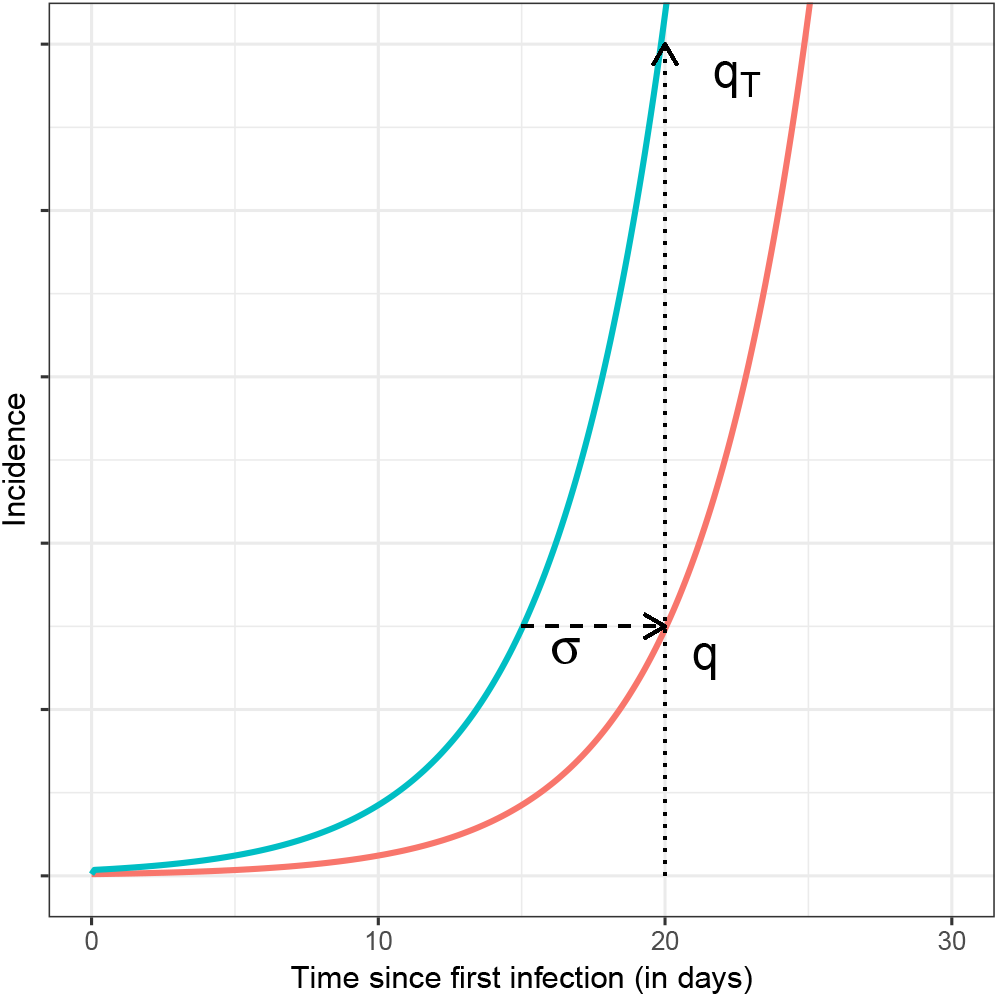
Effect of the asymptomatic period on the incidence. Here this period (*σ* = 5 days) makes a great difference between the detectable incidence *q* and the total incidence *q_T_*.

Following the exponential model, the relation between the total incidence *q_T_* and the detectable incidence *q* is given by:

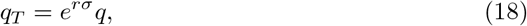

with *σ* the duration of the asymptomatic period. Unlike the exact solution, the approximation smoothly integrates this new epidemic trait. Eq. (15) and (16) become:

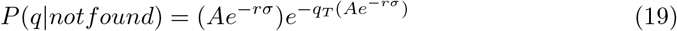

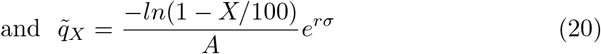

### Application to three pathosystems

Our first example is citrus canker (caused by *Xanthomonas axonopodis* pv. *citri*). Citrus canker can lead to severe losses in commercial citrus (Gottwald et al., 2002). This pathogen has received considerable attention of plant pathology modellers (Parnell et al., 2009; Potts et al., 2013; Neri et al., 2014). It causes lesions on the citrus fruits, stems and leaves, which are diagnostic of pathogen presence during visual inspections. Parameter ranges from literature are reported in Table 1.

**Table 1.** Parameter ranges for the three pathosystem examples.

|  | Minimum | Maximum | Unit | Reference |
| --- | --- | --- | --- | --- |
| Citrus Canker |  |  |  |  |
| $r$ | 0.0155 | 0.0212 | day<sup>-1</sup> | Gottwald et al. (1989) |
| $\sigma$ | 7 | 117 | day | Vernière et al. (2003) |
| Rice Blight |  |  |  |  |
| $r_L$ | $4.4 \cdot 10^{-4}$ | $1.2 \cdot 10^{-3}$ | day<sup>-1</sup> | Mew et al. (1992) |
| $\sigma^\dagger$ | 0.01 | 0.05 | day | see subscript |
| Grey Mould |  |  |  |  |
| $r$ | $8.76 \cdot 10^{-4}$ | $4.57 \cdot 10^{-3}$ | day<sup>-1</sup> | Leroch et al. (2011) |
|  |  |  |  | Angelini et al. (2014) |
|  |  |  |  | Esterio et al. (2017) |
|  |  |  |  | authors data |
| $\sigma$ | 0 | 200 | day | Mcclellan et al. (1973) |
|  |  |  |  | Nair et al. (1995) |
|  |  |  |  | Barnes & Shaw (2002) |

<sup>†</sup> Bacterial blight causes rice tillers to turn yellowish. Fields are inspected from outside, then more closely if looking suspicious. Detection then occurs if infection reaches a threshold incidence $q_d$ . This defines the asymptomatic period as $\sigma = \frac{1}{r_F} \ln \left( \frac{q_d}{q_m} \right)$ , the time lag between strain emergence at individual scale from mutation (at incidence $q_m$ ), with $r_F$ the within field growth rate. Data from Adhikari et al. (1994, 1999).

-2in0in

Our second example, bacterial blight of rice (caused by *Xanthomonas oryzae* pv. *oryzae*), is a serious threat to food security across the globe (Reddy, 1989; Dewa et al., 2011). Breeders have introduced resistance genes into rice cultivars making them absolutely resistant to bacterial blight. However the bacterial species can overcome the resistance and evolve virulent strains. Monitoring programs to establish the absence of virulent strains and/or for early detection of emerging virulence are under development. Observations are done at the field level (rather than at the host level), usually from the roadside. Therefore the relevant *r* value to use in the monitoring model is the landscape scale growth rate (infection from field to field, noted *r_L_*), rather than the within field one (infection from host to host, noted *r_F_*). The parameters values for virulent strains and an explanation of *σ* for this case are given in Table 1 and its subscript.

Our third example concerns grey mould (caused by *Botrytis cinerea*) a fungal plant pathogen of grape (and countless many other hosts). The disease is controlled by fungicide applications but the pathogen can evolve strains less sensitive or insensitive to the fungicide. We consider here the case of insensitivity to Boscalid (a succinate dehydrogenase inhibitor) to which resistance developed in Europe, Australia, the US and South America. Monitoring consists of visits to a large number of grape fields and sampling infections from host individual. Parameter ranges from literature are reported in Table 1.

Although apparently very different in monitoring scale, the case study pathogens can be reported on the same parameter space. Figure 6 locates the epidemics according to their estimated parameters *r* and *σ*. The black crosses figure for each pathogen the likely parameter values, as well as their uncertainty (*i.e.* a long segment shows high variability of the parameter in our sources). If we want to ensure (with 95% confidence) a maximum incidence of 5%, the dashed black contour guides the selection of adequate monitoring effort. Following this curve, we see that this maximum risk can be ensured for the bacterial blight (BB) with only *N* = 20 fields sampled every ∆ = 180 days. The grey mould (GM) case needs a little more frequent monitoring rounds and/or hosts sampled. On the other hand, citrus canker will require *N* = 100 trees to be sampled every ∆ = 30 days, a significantly larger effort.

**Figure 6.**
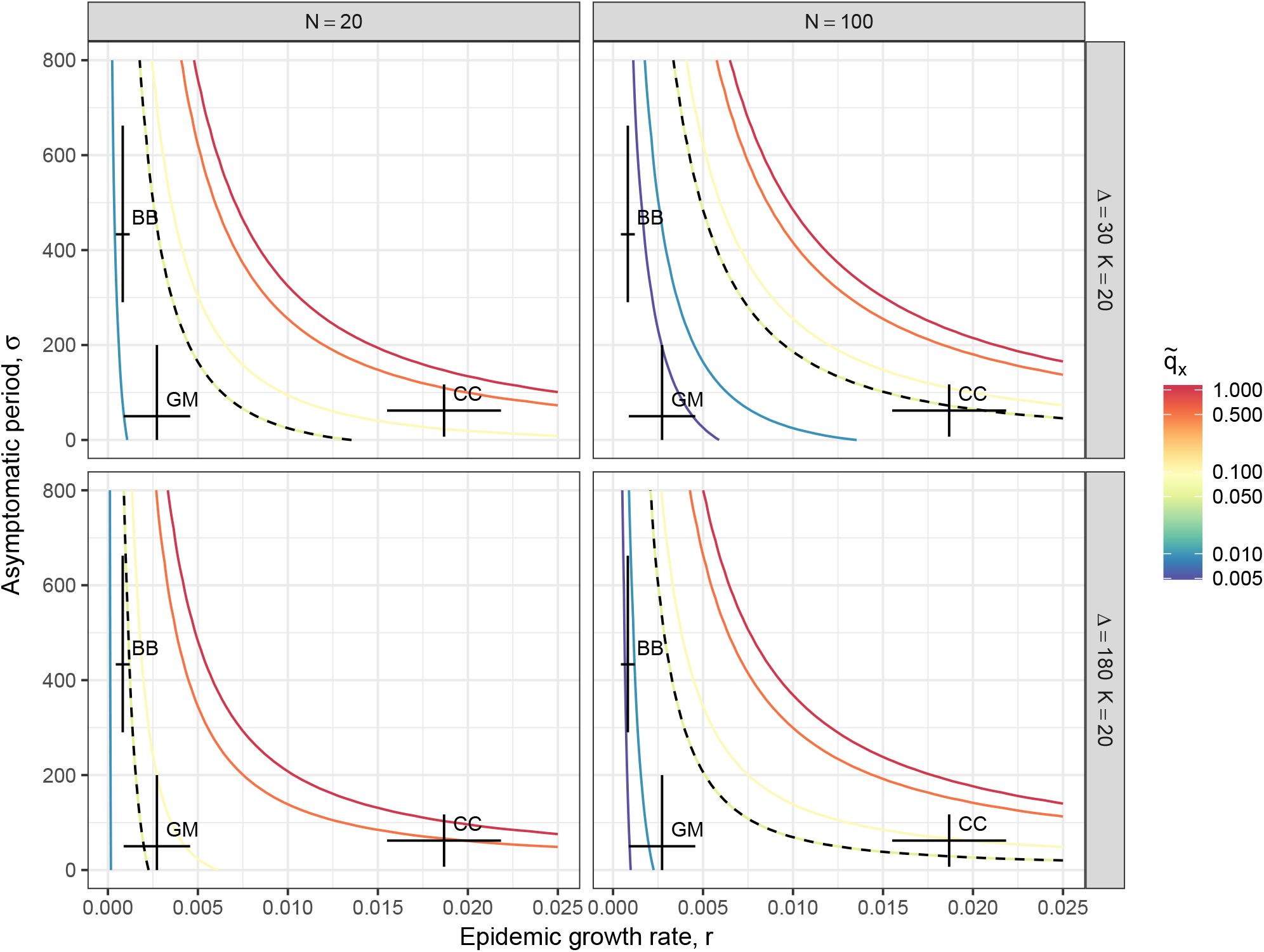
Upper bound of the 95% confidence interval of the incidence, *q*_95_, depending on the epidemic growth rate *r*, and the asymptomatic period *σ*. The contours show the incidence levels whose values are reported on the the log-transformed colour scale bar on the right. The dashed lines figures the 5% maximum incidence. The black crosses figure the likely parameter ranges for citrus canker (CC), bacterial blight (BB) and grey mould (GM).

An interesting output of Figure 6 is the impact of parameter uncertainty on the predicted incidence *q*_95_. For example, although the uncertainty in the *σ* parameter of bacterial blight (BB) is substantial, it is of least concern because it is tangent to the incidence slope (*i.e.* parallel to the contours). However, such level of uncertainty in the *σ* of citrus canker would have cause the “CC black cross” to intersect with all the contour lines, hence predicting a very wide and uninformative range of incidence *q*_95_. In this way, we can quickly assess how input uncertainty will affect the model output, and where more meticulous parameter estimations are required.

These three examples show that, with a combination of crude parameter estimations and a simple calculation, its is possible to assess the monitoring frequency, ∆, and the number of hosts to assess per round, *N*, that are necessary to establish the absence of a pathogen.

## Discussion

The main course of action for infectious disease management resides in monitoring and appropriate response to its outcome. An efficient disease management limits the wasteful use of pesticides, hence reducing their environmental and health consequences while securing their long-term efficiencies. Well-timed responses can also limit the unnecessary culling of hosts (Carpenter et al., 2011; Cunniffe et al., 2015). In addition, monitoring also benefits industries by enabling the certification of pathogen absence which is a primary requirement in the trade of plant and animal produce.

Whether a species is absent or merely undetected is a recurrent question in ecology (Mackenzie, 2005; Wintle et al., 2012). When it comes to pathogens, absence sampling has been addressed according to epidemics specificities, notably with careful attention to the spatial structure of the host populations (Cameron & Baldock, 1998; Coulston et al., 2008). As Metz et al. (1983) did when evaluating the epidemic risk associated with sampling efforts, we account for the epidemic progress between monitoring rounds in our incidence estimation model. Such consideration is essential as we have shown here that assuming a constant incidence over the whole monitoring period leads to severe underestimations of the epidemic progress.

That an epidemiologically informed monitoring proves itself superior to a purely statistical tool like the rule of three is no surprise. Simulation-based approaches are often thoroughly fed with epidemiological knowledge and, so being, have been able to shed light on various aspects of specific diseases like *e.g.* optimal culling ranges (Bates et al., 2003a,b) or economic impacts (Carpenter et al., 2011) for the foot-and-mouth disease. However, such highly specific solutions are not readily valuable for distinct problems. Practical use requires generic tools that are easily accessible and can be straightforwardly applied to observations. Here we propose such a tool in the form of a simple formula, our approximation, which relates a sampling effort to two critical epidemic traits in the form of parameters, namely the growth rate and the asymptomatic period. A subsequent interesting property of our model is that the derived sampling effort can be decomposed in terms of *N*, *K* and ∆, and hence achieved with diverse programs.

It is worth keeping in mind that epidemiologically informed approaches are constrained by the accuracy of the epidemic parameter estimates (Hyatt-Twynam et al., 2017). If our objective is to predict the outcome of an ongoing disease outbreak, parameter estimation must closely follow the detection events, which is often impractical (see *e.g.* Neri et al., 2014). On the other hand, when sampling for disease absence, no observation of the ongoing epidemic exists yet. Parameter estimation is therefore taken from previous occurrences of the epidemic, and possibly from different areas with different environments, or even different hosts species. Occasionally, parameter estimation might also be attempted from outbreaks of a similar disease. Obviously, the cost of widening the origins of observations is an increasing uncertainty on the model outputs. It is then imperative to assess whether or not very crude parameter estimates are acceptable. This can be done conveniently with representations like Figure 6.

Our estimation model relies on the strong hypothesis of the uniformity of *p*(*q*): all incidences are equally likely to be found in the population. More precisely, in our case, the uniformity of *p*(*q*) is ensured at the time of estimation, *i.e.* at the last sampling round. A more common approach consists in ensuring *p*(*q*) uniformity at the first sampling round, calculating a posterior distribution using Bayes’ rule and then using that posterior as prior for the second sample, etc. However, this would not lead to a simple explicit equation like the one we provide, hence limiting its practical use. In both cases, the uniformity of *p*(*q*) seems a bold assumption, as we know that low level incidences are more commonly encountered during monitoring. Nonetheless, assuming uniform *p*(*q*) is a conservative choice, as it biases the estimation towards the safest side: the overestimation of the disease progress.

The model we present here is informed by the temporal dynamics of epidemics. Whether it remains accurate when space becomes part of the system is not obvious, and at some point is likely to depend on host spatial distribution. For example if hosts are clustered in fields, the pathogen dispersion scale and the distance between fields will determine whether or not an epidemic complies with the logistic model underlying this study. Consequently, a direct comparison of our analytical results to spatially explicit simulations should be conducted. In such numerical experiments not only the epidemic but also the sampling process becomes spatially structured, hence breaking the assumption of independent sampling. The robustness of our model in these conditions would therefore be a solid confirmation of its practical value.

## Conclusion

Non-detection is a possible outcome of monitoring programs, but it is an informative one and it can be rendered into a robust risk assessment. Our approximation provides a simple but reliable estimation of pathogen incidence given a sampling effort. It can also be used to derive an appropriate monitoring program for a pathogen, providing that epidemic traits are coarsely known. As it directly builds on elementary parameters of monitoring and epidemic models, this tool can be intuitively adapted to diverse situations as shown by our three examples.

## Acknowledgements

The work at Rothamsted forms part of the Smart Crop Protection (SCP) strategic programme (BBS/OS/CP/000001) funded through the Biotechnology and Biological Sciences Research Council’s Industrial Strategy Challenge Fund. Authors are also thankful to the Newton Fund of the British Council, the US Department of Agriculture as well as the National Science and Technology Development Agency (NSTDA) of Thailand, Curtin University, the Grains Research and Development Corporation (GRDC) as well as the Australian Grape and Wine Authority for funding support. Authors wish to thank Mr Lincoln Harper for providing fungicide resistance data on grey mould.

## Footnotes

1 We use here the plant pathology definition where virulence signifies the ability of the pathogen to infect the host. In human and other animal pathology virulence is used as a measure of damage the pathogen does to the host.

2 We use here the plant pathology definition where incidence is the fraction of host units infected. In human and other animal pathology this is termed prevalence.

